# Yeast single cell protein production from a biogas co-digestion substrate

**DOI:** 10.1101/766345

**Authors:** Jonas A. Ohlsson, Matilda Olstorpe, Volkmar Passoth, Su-lin L. Leong

**Affiliations:** Department of Molecular Sciences, Swedish University of Agricultural Sciences, Uppsala, Sweden

## Abstract

Biogas plants serve as hubs for the collection and utilization of highly nutritious waste streams from households and agriculture. However, their outputs (biogas and digestate) are of relatively low economic value. Here, we explore the co-production of yeast single cell protein, a potentially valuable feed ingredient for aquaculture and other animal producing industries, with biogas on substrate collected at a co-digestion biogas plant, using three yeast species well suited for this purpose (*Wickerhamomyces anomalus*, *Pichia kudriavzevii*, and *Blastobotrys adeninivorans*). All yeasts grew rapidly on the substrate, yielding 7.0–14.8 g l^−1^ biomass after 12–15 The biomass crude protein contents were 22.6–32.7 %, with relatively favorable amino acid compositions mostly deficient in methionine and cysteine. Downstream biomethanation potential was significantly different between yeast species, with the highest product yielding species (*Blastobotrys adeninivorans*) also yielding the highest biomethanation potential.

**Highlights:**

- All yeasts grew well on the biogas substrate, with high growth rates.
- Produced biomass was of high nutritional value for use in fish feed formulations.
- Downstream effects on methane potential were strain-dependent.
- Yeast biomass may be a viable biogas co-product.

## 1 Introduction

The organic fraction of municipal solid waste (OFMSW) is a major waste stream which contains ample amounts of energy and nutrients available for microbial conversion into higher value products. In the EU, policy changes such as the EC Council Directive 1999/31/EC of 26 April 1999, by which member states are obligated to reduce the amount of OFMSW being deposited on landfills, have increased demands for alternative waste processing facilities such as anaerobic digestion (AD) plants. AD plants convert waste streams into biogas and a post-digestion residue mainly being used as fertilizer. OFMSW is typically co-digested with agricultural residues such as manure, a practice which improves biogas yields as well as reduces emissions of methane and odours from farms (Holm-Nielsen et al., 2009).

Moreover, biogas production by AD is in line with other policies aimed at improving energy security and usage of renewable energy. In concert with the implementation of several such policies, the installed capacity of AD in Europe has seen a steep increase since the turn of the century (Scarlat et al., 2018), and future demand may rise further if new uses such as biomethane-based power grid balancing, which offsets energy fluctuations due to large parts of solar and wind in the energy mix, see widespread deployment (Peters et al., 2018).

The widespread adoption of OFMSW as a substrate for AD has at least two main implications that warrant considering AD plants as well suited for expanding their product portfolios: first, the year-round and constant availability of readily biodegradable substrate of relatively stable composition (Hartmann and Ahring, 2006), and, second, the presence of well-developed logistic networks for the collection and transportation of substrate. Furthermore, AD substrate is typically pretreated in several ways including removal of certain contaminants, size reduction, and hygienization; unit operations which render the material more amenable to microbial conversion (Carlsson et al., 2012).

Although biogas and digestate represent outputs that provide value, both are low-value commodities. There are several recently proposed methods for improving the financial proposition of biogas plants by altering the output streams towards more valuable products (see e.g., Kleerebezem et al. (2015), Monlau et al. (2015)). One possible value-added co-product of a biorefinery is yeast biomass intended as animal feed. A particularly promising consumer of microbial protein feed sources is the aquaculture industry. While outputs from marine fisheries are probably declining (FAO, 2018), aquaculture today supplies 53 % of direct global fish consumption (FAO, 2018), and as the fastest growing food producing sector over the last 40 years it will play a crucial role in feeding the growing world population (Béné et al., 2015). A major ingredient in aquaculture feed, especially for carnivorous fish species is fish meal (FM), which is derived from wild-caught fish. With an ever-widening gap between production of wild and farmed fish, availability of FM is becoming a major limiting factor for further expansion of fish farming (Tacon and Metian, 2009). Although feed ingredients of vegetable origin are used for replacing FM in feed formulations, higher inclusion levels may negatively affect fish growth due to antinutritional factors present in many vegetable feed sources (Francis et al., 2001). Moreover, vegetable feed ingredients, as well as FM, may directly be at odds with human food interests (Tacon and Metian, 2009).

Microbial biomass used for feed is commonly referred to as single cell protein (SCP), and its use in animal feed is well established for both fish and other animals (Goldberg, 1985). Targeting fish feed is especially beneficial as fish are able to metabolize the nucleic acids that are present in large amounts in yeast biomass, whereas high contents of nucleic acids in the diet can cause adverse health effects in some other animals (Rumsey et al., 1992). Yeast biomass has been successfully used in fish feed formulations, both at low inclusion levels for its appetite stimulating, nutritional, or immunostimulatory properties, and at higher levels as FM replacement where it is generally well tolerated up to a certain level (reviewed in Delgado and Reyes (2018)). SCP production is desirable from a food security perspective, and the demand for fish feed ingredients, including FM replacements, are likely to increase due to the concomitant expansion of aquaculture and decrease in marine fish stocks.

Organisms suitable for SCP production on OFMSW should, first and foremost, be able to utilize a large array of substrate molecules. Other desirable characteristics include phytase production, as this may improve nutritional quality of the feed if ingredients of vegetable origin are included in the formulation (Cao et al., 2007), as well as the ability to outcompete other organisms due to the non-sterile nature of the substrate. In this study, we evaluated three yeast species with suitable properties for SCP production on typical co-digestion AD substrate: *Wickerhamomyces anomalus*, a metabolically versatile species which has shown robustness to difficult growth conditions, has been evaluated in fish feeding trials and which is known for its biocontrol properties as well as being a phytase producer (Huyben et al., 2017; Olstorpe et al., 2009; Passoth et al., 2006; Passoth et al., 2010; Schnürer and Jonsson, 2010); *Pichia kudriavzevii*, another robust yeast known for its ability to grow in the presence of inhibitory substances, and which has been reported to produce extracellular phytase (Hellström et al., 2015; Olstorpe et al., 2009; Radecka et al., 2015), and *Blastobotrys adeninivorans*, known for its ability to utilize a large variety of carbon and nitrogen sources, as well as for its high production of intra- and extracellular phytase (Middelhoven et al., 1991; Olstorpe et al., 2009; Sano et al., 1999).

The aim of this study was to investigate whether production of yeast biomass in combination with AD could be a feasible option for further diversification of production outputs from biogas plants. We have evaluated growth performance of yeasts on biogas substrate obtained from a Swedish co-digestion biogas plant, characteristics of the resulting biomass, and the effects of yeast cultivation on downstream chemical composition and biomethanation potential (BMP).

## 2 Materials and methods

### 2.1 Inoculum preparation and culture media

Yeast strains (*W. anomalus* CBS 100487, *P. kudriavzevii* CBS 2062, and *B. adeninivorans* CBS 7377), stored in 50% glycerol stocks at −80°C, were inoculated onto YPD agar (10 g l^−1^ yeast extract (BD, Le Pont-de-Claix, France), 20 g l^−1^ bacterial peptone (BD, Le Pont-de-Claix, France), 20 g l^−1^ D-glucose (Merck, Darmstadt, Germany), and 20 g l^−1^agar (BD, Le Pont-de-Claix, France)). Inoculum cultures were prepared using the same medium, without agar, in 125-ml baffled Erlenmeyer flasks (Thomson Ultra-Yield, Thomson Instrument Co., Carlsbad, CA, USA), and cultivated on a rotary shaker for 24 h. Cells were harvested at 3000 × *g* for 5 min and washed with saline (NaCl, 9 g l^−1^) using the same settings.

### 2.2 Substrate preparation

The biogas substrate was obtained directly from the inlet to the digester at a biogas plant in Sweden, and consisted mainly of source-separated household waste, organic waste from municipal kitchens, and liquid agricultural waste (swine and cattle manure). Metals and plastics had been mechanically removed at the biogas plant, and the substrate had been hygienized at 70°C for >1 h. This substrate will be referred to as *native substrate (NS)*.

To be able to separate yeast biomass after culturing, and to reduce the risk of contamination as substrate was collected through a non-sterile sampling port at the biogas plant, the substrate was sterile-filtered. This was accomplished using an Asahi Rexeed-25A hemodialyzer (Asahi Kasei Medical Co., Ltd., Tokyo, Japan) connected to a peristaltic pump, with the filter replaced when the counterpressure reached 0.6 bar. The filtered substrate was then sterile-filtered through a 0.2 μm sodium acetate filter (Nalgene Rapid-Flow, Thermo Fisher Scientific, Waltham, MA, USA) using a Büchner funnel.

### 2.3 Bioreactor operation

500-ml Infors HT Multifors CSTR bioreactors (Infors AG, Bottmingen, Switzerland) were used for the cultivations. For each reactor, 400 ml of the sterile-filtered substrate was inoculated at an initial OD_600_ of 1.0. Reactor parameters were pH = 7.00 ± 0.10, stirrer = 500 rpm, and pO_2_ = 0.2. pO_2_ was maintained using stirrer speed, with a minimum of 200 rpm and a maximum of 1200 rpm. pH was automatically adjusted on-line using 5 M NaOH and 3 M H_3_PO_4_.

Fermenter temperature was set to 30°C for all cultivations, except for the evaluation of *B. adeninivorans* CBS 7377 growth performance. This strain exhibited better growth at 37°C during initial growth assessment (results not shown), so growth was evaluated at this temperature

It was not possible to maintain a pO_2_ value of 0.2 during the exponential growth phase, due to the high oxygen consumption of the yeast. Cultivations were terminated after the log phase was completed, as indicated by pO_2_ readings.

To monitor the fermentations, viable cell counts were performed. Due to the complexity and optical activity of the medium, plating was chosen instead of OD measurements. Relevant dilutions, made with 1 g l^−1^ peptone water, were plated onto YMC agar (3 g l^−1^ yeast extract, 3 g l^−1^ malt extract (BD, Le Pont-de-Claix, France), 5 g l^−1^ bacterial peptone, 10 g l^−1^ D-glucose, 100 mg l^−1^ chloramphenicol (Sigma-Aldrich, Steinheim, Germany)). Plates were incubated at 30°C and colony forming units (CFU) were counted when colonies were clearly visible.

After fermentation, bioreactor contents were centrifuged at 3000 × *g* for 10 min. The pellets, containing yeast biomass, were washed with deionized water using the same centrifuge settings, and stored at −20°C.

### 2.4 Biomethanation potential assay

To assess downstream effects of yeast cultivation on biogas performance, spent medium was collected from cultivations of each yeast. Cultivations were performed largely as described in Section 2.3. In order to minimize confounding factors, all cultivations were terminated at the same time (i.e., the time was determined by the growth performance of the slowest growing yeast), and were run at the same temperature (30°C). This was needed to ensure that evaporative losses of volatile energy carriers, such as short-chain carboxylic acids, were similar between the treatments. At the end of cultivation, yeast biomass was collected as described in Section 2.3, and the supernatants, referred to as *spent media*, collected.

The BMP assay was conducted largely according to Angelidaki et al. (2009). In brief, total solids (TS) and volatile solids (VS) of NS and spent medium (supernatants, post yeast-treatment) were determined by drying the substrates at 105°C and incinerating at 550°C in aluminum containers, noting the weights after each step. TS was calculated as the quotient of dry matter divided by initial weight. VS was determined as the difference between TS and ash content.

The assay was performed using untreated NS and fresh inoculum (collected from the same biogas plant and degassed for 3 days at 37°C), contributing approximately 1.2 g VS and 3.6 g VS, respectively. The substrate control treatment contained only NS and inoculum. Spent medium was added in the remaining treatments, so that the mixtures contained, by weight, NS:spent medium in ratios of 10:1, 10:3, 2:1, and 1:1, which corresponded to 9–50% spent medium in the final AD slurry. Due to the low VS content of the spent medium, increases in VS due to supernatant additions were modest, at most 20%, and it was assumed that this slight change in inoculum:substrate ratio would not affect inoculum performance. Inoculum and cellulose process controls were included. The inoculum control, used for determination of background methane production, consisted of inoculum contributing 3.6 g VS. The cellulose control, needed for evaluating the function of the inoculum, contained 3 g cellulose (medium fibers; Sigma-Aldrich, Steinheim, Germany) and 400 ml of inoculum. Tap water was added to each bottle to a final volume of 400 ml, and each treatment was evaluated in triplicate.

To measure methane production, AMPTS II Automatic Methane Potential Test Systems (BioProcess Control AB, Lund, Sweden) were used, detailed in Badshah et al. (2012). Briefly, gas volume measuring was performed using a piston system, and gas was upgraded by flushing it through 7 M NaOH. Samples were agitated using stirrers attached directly to the bottles. The assay was performed at 37°C, as this was the temperature used in the commercial digester from which the inoculum was derived. Bottles were flushed with N_2_ gas prior to initiating the experiment. Specific methane potential was obtained by dividing the volume of methane produced by the actual amount of VS in each sample. When less than one reading per day was generated from all samples (i.e. the piston did not register any methane gas emissions for 24 h), the assay was determined to be complete. Each AMPTS II system contained an identical set of samples so that the response of each system could be included in the statistical model (Section 2.6).

### 2.5 Chemical analyses

Chemical analyses were purchased from external labs. Crude protein (CP) content was determined using the total nitrogen Kjeldahl method, and CP was calculated as N × 6.25 (Nordic Committee on Food Analysis, 2003). Crude lipid (CL) content was determined according to (The Commission of the European Communities, 1998). Gross nutritional analyses were performed at the VHC lab (SLU, Uppsala, Sweden). Amino acid (AA) analyses were performed according to the ISO 13903:2005 method (Eurofins Food & Agro, Jönköping, Sweden). Micronutrient compositions of spent medium (supernatants, yeast biomass removed), sterile-filtered medium, and NS were analyzed using ICP-MS (Agrilab AB, Uppsala, Sweden). Dry matter was determined by drying the samples at 105°C until constant weight was achieved.

When possible, analyses were performed in replicates. For the BMP assay, each yeast treatment was carried out in a single bioreactor to ensure equal freshness and composition of the spent medium, and the replicates represent aliquots of the supernatants. For pellet (yeast biomass) characterization, four replicate fermentations were carried out. Spent medium compositional analyses were performed using material pooled from all bioreactors.

### 2.6 Calculations and statistical analyses

The maximum growth rate (μ_max_) was estimated by taking the greatest slope of the growth curve

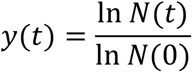

where *N(t)* is the number of CFU at timepoint *t*. Final biomass productivity was calculated by dividing final biomass concentration by the total cultivation time. Maximum growth rates and final productivities were calculated independently for each fermenter.

AA scores for the yeast biomass, reflecting the requirements of essential amino acids (EAA), were calculated based on finfish requirements as reported in Tacon et al. (2009).

To evaluate the downstream effects of spent substrate addition on biomethanation potential, a linear model was fitted according to the formula

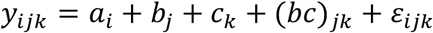

where *y_ijk_* is the biomethanation potential of the sample, *a* is the effect of each AMPTS II system, *b* is the effect of strain, *c* is the effect of spent medium at level *k* in the reactor, *(bc)* is the interaction term for dose × strain, and *ε* is the residual. The model was fitted using the built-in function lm() in R version 3.6.0 (R Core Team, 2019). All graphs were generated using ggplot2 version 3.1.1 (Wickham, 2009).

## 3 Results

### 3.1 Yeast cultivation and biomass characterization

Native biogas substrate (NS) was sterile-filtered and the three yeast strains were cultured on this substrate in the fermenters. Fermentations were terminated after 12–15 hours, yielding approximately 2 log_10_ increases for strains *P. kudriavzevii* CBS 2062 and *B. adeninivorans* CBS 7377, and approximately a 1.5 log_10_ increase for *W. anomalus* CBS 100487 (Figure 1), with high maximum growth rates (μ_max_) of 0.48–0.76 h^−1^ (Table 1).

**Figure 1.**
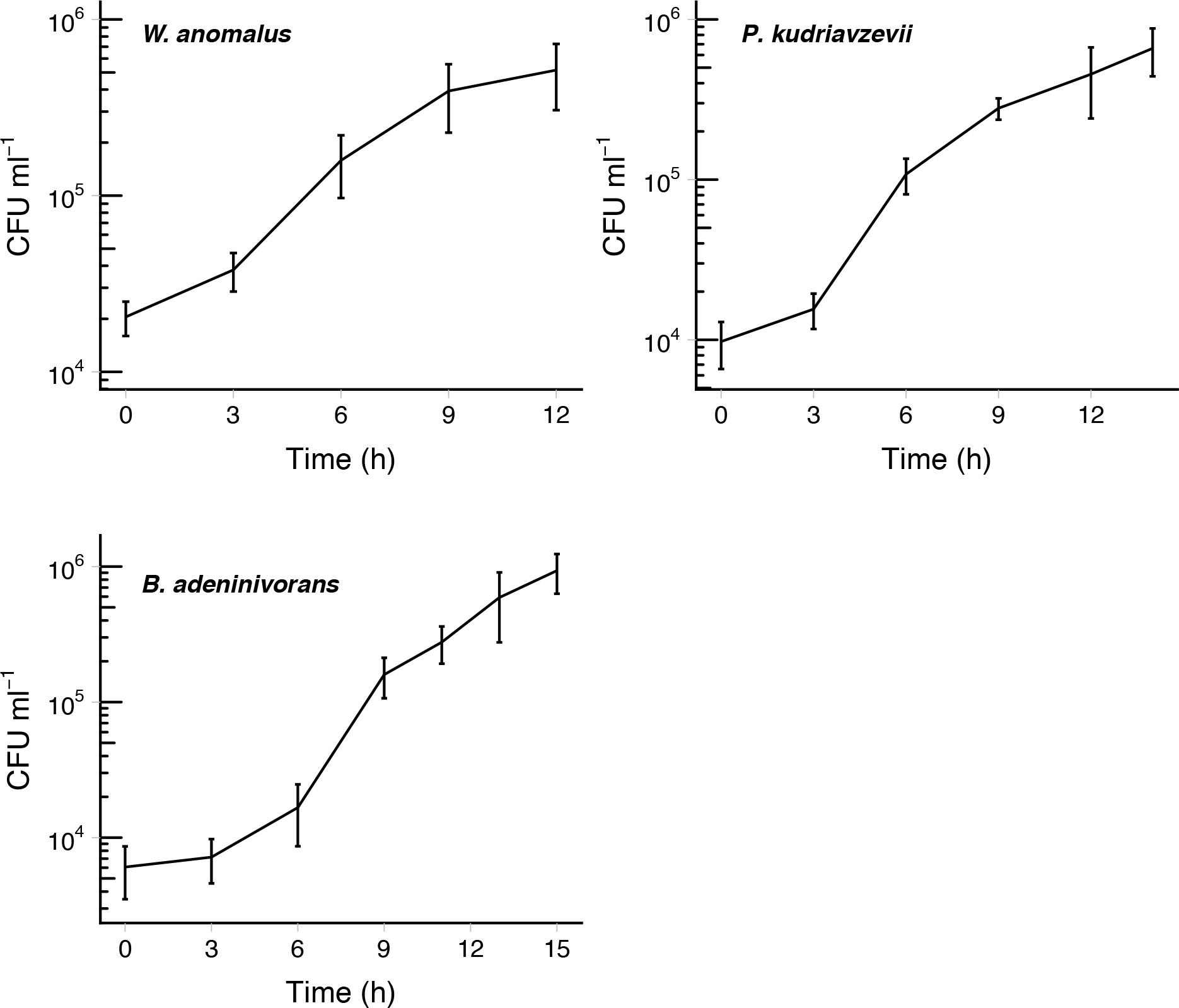
Growth curves for the yeast cultivation trials. Bars represent standard deviation. *W. anomalus* and *P. kudriavzevii* were grown at 30°C, and *B. adeninivorans* at 37°C.

**Table 1.** Nutritional characterization of washed yeast biomass, on a dry matter basis. Missing standard deviations indicate that measurements were performed on pooled replicates

| Parameter | *W. anomalous* |  | *P. kudriavzevii* |  | *B. adeninivorans* |  |
| --- | --- | --- | --- | --- | --- | --- |
|  | Value | SD | Value | SD | Value | SD |
| *N* | 3<sup>a</sup> |  | 4 |  | 3<sup>a</sup> |  |
| *Biomass concentration (g/l)* | 7.03 | 0.08 | 7.36 | 0.34 | 14.83 | 1.02 |
| *μ<sub>max</sub> (h<sup>-1</sup>)* | 0.48 | 0.10 | 0.64 | 0.06 | 0.76 | 0.06 |
| *Final biomass productivity (g/l·h)* | 0.59 | 0.01 | 0.53 | 0.02 | 0.99 | 0.07 |
| *Crude energy (MJ/kg)* | 11.05 | – | 11.91 | – | 13.05 | – |
| *Crude protein (%)* | 22.59 | 0.35 | 32.71 | 0.54 | 30.46 | 1.22 |
| *Ash (%)* | 45.77 | 0.42 | 42.88 | 0.85 | 37.58 | 2.92 |
| *Crude lipid (%)* | 2.68 | – | 1.76 | – | 1.56 | – |
| *Ca (g/kg)* | 145.6 | – | 123.6 | – | 115.0 | 7.23 |
| *K (g/kg)* | 7.4 | – | 11.1 | – | 16.4 | 2.07 |
| *P (g/kg)* | 6.6 | – | 11.1 | – | 6.9 | 2.57 |
| *Mg (g/kg)* | 4.7 | – | 3.7 | – | 11.2 | 0.80 |
| *Na (g/kg)* | 83.3 | – | 85.3 | – | 76.7 | 0.93 |
| *S (g/kg)* | 3.0 | – | 3.6 | – | 4.3 | 0.40 |

<sup>a</sup> One sample was excluded due to equipment malfunction.

Final biomass concentrations ranged from 7.0 g l^−1^ (22.6% CP) for *W. anomalus* to 14.8 g l^−1^ (30.5% CP) for *B. adeninivorans*. Final biomass productivities ranged from 0.53 g l^−1^ h^−1^ for *P. kudriavzevii* to 0.99 g l^−1^ h^−1^ for *B. adeninivorans* (see Table 1 for the kinetic parameters and gross nutritional characterization of the yeast biomass).

Protein content according to the AA analysis was largely in agreement with CP values, indicating that most nitrogen present in the biomass originated from protein (Table 2). Yeast biomass AA composition was similar for all three species, with *W. anomalus* notably having a somewhat lower content of methionine and cysteine, sulfur-containing EAA, compared to the other two yeasts. In general, yeast biomass was deficient in arginine and in the sulfur-containing AA relative to the requirements of finfish, whereas other AA where either in excess or at similar levels (Table 2). EAA made up approximately half of the AA present in yeast biomass.

**Table 2.** Amino acid composition of dried yeast biomass. Tryptophan contents were not analyzed

|  | *W. anomalus* |  |  | *P. kudriavzevii* |  |  | *B. adeninivorans* |  |  |
| --- | --- | --- | --- | --- | --- | --- | --- | --- | --- |
|  | Content<br>(g/100 g) | Content<br>(weight<br>%) | AA<br>score<sup>a</sup> | Content<br>(g/100 g) | Content<br>(weight<br>%) | AA<br>score | Content<br>(g/100 g) | Content<br>(weight<br>%) | AA<br>score |
| *Alanine* | 1.55 | 8.8 |  | 1.71 | 6.4 |  | 1.93 | 7.7 |  |
| *Arginine* | 0.87 | 5.0 | 85 | 1.35 | 5.0 | 82 | 1.36 | 5.4 | 95 |
| *Aspartic acid* | 1.88 | 10.7 |  | 3.32 | 12.4 |  | 2.42 | 9.7 |  |
| *Cysteine* | 0.157 | 0.9 | 67 | 0.274 | 1.0 | 70 | 0.275 | 1.1 | 81 |
| *Glutamic acid* | 2.63 | 15.0 |  | 3.73 | 13.9 |  | 4.08 | 16.4 |  |
| *Glycine* | 0.84 | 4.8 |  | 1.30 | 4.9 |  | 1.28 | 5.1 |  |
| *Histidine* | 0.351 | 2.0 | 83 | 0.594 | 2.2 | 88 | 0.572 | 2.3 | 96 |
| *Hydroxyproline* | <0.05 | — |  | <0.05 | — |  | <0.05 | — |  |
| *Isoleucine* | 0.87 | 4.9 | 131 | 1.42 | 5.3 | 133 | 1.1 | 4.4 | 120 |
| *Leucine* | 1.36 | 7.7 | 114 | 2.11 | 7.9 | 110 | 1.84 | 7.4 | 110 |
| *Lysine* | 1.41 | 8.0 | 95 | 2.22 | 8.3 | 93 | 1.74 | 7.0 | 84 |
| *Methionine* | 0.242 | 1.4 | 50 | 0.529 | 2.0 | 69 | 0.417 | 1.7 | 63 |
| *Ornithine* | <0.01 | — |  | <0.01 | — |  | <0.01 | — |  |
| *Phenylalanine* | 0.84 | 4.8 | 100 | 1.31 | 4.9 | 97 | 1.09 | 4.4 | 94 |
| *Proline* | 0.75 | 4.3 |  | 1.06 | 3.9 |  | 1.44 | 5.8 |  |
| *Serine* | 1.06 | 6.1 |  | 1.49 | 5.6 |  | 1.44 | 5.8 |  |
| *Threonine* | 0.99 | 5.6 | 106 | 1.66 | 6.2 | 110 | 1.39 | 5.6 | 107 |
| *Tyrosine* | 0.76 | 4.4 | 134 | 1.15 | 4.3 | 125 | 1.00 | 4.0 | 125 |
| *Valine* | 0.96 | 5.5 | 115 | 1.58 | 5.9 | 117 | 1.55 | 6.2 | 132 |
| *Total* | 17.52 | 100.0 |  | 26.81 | 100.0 |  | 24.92 | 100.0 |  |
| *Total EAA* | 8.81 | 50.3 |  | 14.20 | 53.0 |  | 12.33 | 43.0 |  |

<sup>a</sup> The AA score reflects dietary requirements of essential amino acids in finfish, with values below 100 indicating deficiency. Essential amino acid requirements of finfish are according to Tacon et al. (2009).

### 3.2 Characterization of spent medium

Filtration of the substrate removed a considerable amount of nutrients, presumably contained in particulate matter. Compared to NS, filtered substrate contained 5-fold less VS, 2.1-fold less total N, and 4.5-fold less total C (Table 3). Compared to the filtered supernatant, spent medium contained less TS, VS, C and N, and generally reduced levels of micronutrients. The exceptions were Na, which increased in all treatments, and P, which increased in two out of three treatments. This is most likely due to the use of NaOH and H_3_PO_4_ for pH control. Treatment with *P. kudriavzevii* yielded the lowest reduction in TS and VS.

**Table 3.** Chemical composition of substrate before (native) and after filtration, and after yeast fermentation

|  | Native | Filtered | *Post-fermentation supernatants* |  |  |
| --- | --- | --- | --- | --- | --- |
|  |  |  | *W. anomalus* | *P. kudriavzevii* | *B. adeninivorans* |
| *TS (%)* | 7.8 | 2.1 | 1.5 | 1.6 | 1.2 |
| *VS (% of TS)* | 86.3 | 67.2 | 49.8 | 63.9 | 47.9 |
| *Total N (g kg<sup>-1</sup>)* | 2.7 | 1.3 | 0.6 | 0.5 | 0.3 |
| *Organic N (g kg<sup>-1</sup>)* | 2.2 | 0.8 | 0.2 | 0.4 | 0.2 |
| *NH<sub>4</sub><sup>+</sup>-N (g kg<sup>-1</sup>)* | 0.5 | 0.4 | 0.4 | 0.2 | 0.1 |
| *Total C (g kg<sup>-1</sup>)* | 36.3 | 8.1 | 5.0 | 5.1 | 3.4 |
| *C/N ratio* | 13.3 | 6.3 | 8.5 | 9.4 | 11.0 |
| *P (g kg<sup>-1</sup>)* | 0.37 | 0.20 | 0.15 | 0.74 | 0.39 |
| *K (g kg<sup>-1</sup>)* | 1.37 | 1.25 | 1.19 | 1.14 | 0.96 |
| *Mg (g kg<sup>-1</sup>)* | 0.24 | 0.19 | 0.08 | 0.06 | 0.09 |
| *Ca (g kg<sup>-1</sup>)* | 1.77 | 1.23 | 0.06 | 0.07 | 0.03 |
| *Na (g kg<sup>-1</sup>)* | 0.52 | 0.51 | 2.09 | 1.27 | 1.70 |
| *S (g kg<sup>-1</sup>)* | 0.24 | 0.08 | 0.05 | 0.06 | 0.04 |

### 3.3 BMP assay

The BMP assay was terminated after 40 days. The methane potential of native substrate with addition of spent substrate was significantly different depending on strain choice (*p* = 0.0009), with no effect of the dose of spent medium added or interaction of strain and dose observed (Table 4). Native substrate with spent medium had BMP levels of 342.0–380.5 ml CH_4_/g VS (Table 5). Native substrate had a BMP of 323.0±17.0 ml CH_4_/g VS. Cellulose controls yielded 392.3±31.0 ml CH_4_/g, indicating that the inoculum performed adequately, and inoculum controls produced 70.7±21.3 ml CH_4_.

**Table 4.** ANOVA table for effects of strain choice and spent medium dosage on downstream biomethanation potential

| **Factor** | **DF** | **Sum sq.** | **Mean sq.** | **F value** | **p-value** |
| --- | --- | --- | --- | --- | --- |
| *AMPTS system* | 2 | 7595.2 | 3797.6 | 10.1089 | 0.0005 |
| *Strain* | 2 | 6890.2 | 3445.1 | 9.1707 | 0.0009 |
| *Dose* | 1 | 14.3 | 14.3 | 0.0381 | 0.8467 |
| *Strain:Dose* | 2 | 2284.8 | 1142.4 | 3.0410 | 0.0644 |
| *Residuals* | 27 | 10143.0 | 375.7 |  |  |

**Table 5.** Biomethanation potential (BMP) of native substrate with addition of spent medium from the yeast cultivations according to the fitted linear model

| **Strain** | **BMP (ml CH<sub>4</sub>/g VS)** | **SE** |
| --- | --- | --- |
| *W. anomalus* | 350.7 | 13.0 |
| *P. kudriavzevii* | 342.0 | 17.5 |
| *B. adenivorans* | 380.5 | 17.2 |

## 4 Discussion

In this study, we have produced yeast biomass from the liquid fraction of a substrate from a Swedish co-digestion biogas plant. Depending on the yeast strain, biomass concentrations reached 7.0–14.8 g/l after 12–15 h. By including the spent medium from yeast fermentation in the AD slurry, we were also able to assess the relative effects of each yeast strain on downstream methane potential.

For yeast-based SCP to be a viable co-product at a biogas plant, the organism must be able to utilize the substrate used at the plant. The species evaluated here grew well on the biogas substrate, with the highest final biomass productivity being 0.99 g l^−1^ h^−1^, with only minor efforts made to optimize cultivation conditions. Initial experiments, carried out in shake flasks and in aerated 96-well deep-well plates, were conducted to determine suitable fermentation parameters, but no subsequent optimization was conducted. Furthermore, fermentations were performed at the same pH for all strains, in order to get representative spent substrates for the BMP assay. It is worth noting that none of the yeast strains were able to grow at a pH of below 6, likely due to the presence of weak acids in the substrate. The substrate otherwise proved non-toxic to the yeast and no dilution was necessary. This is important for a biogas plant, as further addition of water would lower the organic loading rate and hydraulic retention time, compromising methane production.

Crude protein contents of the yeast biomass were lower than what has been reported in the literature for fodder yeast, which range from 34.3–48.2 on a dry weight basis (Tacon et al., 2009), whereas the levels in this study ranged from 22.6 to 32.7. However, much of the crude protein was accounted for by the amino acid analysis, indicating low levels of non-protein nitrogen sources. Amino acid compositions of the yeast biomass were similar to those found in the literature for other yeasts used as fish feed (Tacon et al., 2009), with methionine being the most limiting amino acid. Notably, lysine levels were close to the requirements of finfish. Lysine is commonly a limiting amino acid, especially in feed ingredients of vegetable origin but also in many species of yeast (Tacon et al., 2009). With further optimization, it is likely that the protein content could be increased. Rajoka et al. (2006), in a study using *Candida utilis*, found that true protein contents increased during the first 24 hours of cultivation. Likewise, for *Saccharomyces cerevisiae* it was shown that protein contents were highest after 36 h, for two out of three strains evaluated (Novak, 2007). In the present study, however, cultivations were terminated after 12–15 h, suggesting an obvious route for optimization.

The choice of yeast strain proved to be important both for product yields as well as for downstream BMP. The best-performing yeast strain, *B. adeninivorans*, produced approximately twice the amount of biomass compared to the other two species, with a similar protein content. Accordingly, final biomass productivity was also approximately double compared to the two other strains. It is possible that this is due in part to the higher cultivation temperature used for this strain. Rajoka et al. (2006) found that temperature had a large impact on crude protein contents of yeast biomass, which they attributed to increased transport of nutrients over the cell membrane. Interestingly, *B. adeninivorans* also had the highest BMP when mixing its spent substrate with native biogas substrate, suggesting a synergistic effect of this yeast treatment on the AD process and highlighting the importance of yeast strain choice for overall financial viability. It is worth mentioning that the production of yeast biomass in all cases exceeded the reduction of TS in the substrate during cultivation. This finding is an artifact of the way TS and VS are measured, which discounts all compounds with a boiling point below 105°C, such as several organic acids abundantly present in household waste.

Although the results of this initial study are promising, future scale-up studies are needed need to address a number of limitations, some of which are summarized below. First, cultivations should be performed on minimally filtered substrate. In this study, sterile-filtered substrate was used in order to facilitate analyses, such as biomass dry matter concentration and CFU counts. Despite the substrate being hygienized (70°C for at least one hour), it cannot be regarded as sterile and it remains to be seen whether this proves a challenge in scaling up. Second, efficient product recovery will be essential for the financial outcome of the process, but was not investigated in the present study. Third, the downstream effects on methane production should be more rigorously characterized. Whereas the present study used batch tests for assessing methane potential, a more thorough study would use a continuous biogas reactor with continuous feeding of spent substrate. Further, although the results suggest an increase in methane potential over the native substrate, it is fair to assume that sterile filtration of the substrate acts as a pretreatment by removing the more recalcitrant particulate matter. Thus, future studies, using the setup described above and including the retentate in the biogas reactor, could better assess the effects of yeast cultivations on the economics of the integrated process, preferably by examining effects on different types of biogas substrates using several inocula.

## 5 Conclusions

Biogas plants are in several ways well suited for production of products other than methane and digestate. In this initial investigative study, we demonstrate that co-production of yeast SCP on biogas co-digestion substrate is feasible, with high growth rates attained during the batch cultivations. Furthermore, yeast biomass amino acid profiles were similar to yeast SCP already in use in the aquaculture industry. Due to the presence of logistics networks and year-round substrate availability, co-production of yeast SCP and biogas may be an attractive option for diversifying biogas plant outputs. Further optimization of cultivation parameters is likely to improve product yield and productivity; however, scale-up experiments are required to assess the financial and technological viability of this integrated process.

## Acknowledgements

The authors wish to thank Simon Isaksson and Albina Bakeeva for help with setting up the BMP assay.

## List of abbreviations

μmax: Maximum growth rate
AA: Amino acid(s)
AD: Anaerobic digestion
BMP: Biomethanation potential
CL: Crude lipid
CP: Crude protein
EAA: Essential amino acid(s)
FM: Fish meal
NS: Nativesubstrate
OFMSW: Organic fraction of municipal solid waste
SCP: Single cell protein
TS: Total solids
VS: Volatile solids

